# Denoising physiological data collected during multi-band, multi-echo EPI sequences

**DOI:** 10.1101/2021.04.01.437293

**Authors:** Katherine L. Bottenhorn, Taylor Salo, Michael C. Riedel, Matthew T. Sutherland, Jennifer L. Robinson, Erica D. Musser, Angela R. Laird

## Abstract

Collecting physiological data during fMRI experiments can improve fMRI data cleaning and contribute to our understanding of psychophysiological processes; however, these recordings are frequently fraught with artifacts from the MRI pulse sequence. Here, we assess data from BIOPAC Systems, Inc., one of the more widely-used manufacturers of physiological monitoring equipment, and evaluate their recommendations for filtering such artifacts from electrocardiogram and electrodermal activity data collected during single-band, single-echo fMRI sequences and extend these recommendations to address artifacts associated with multiband, multi-echo fMRI sequences. While the magnitude and frequencies of artifacts differ with these aspects of pulse sequences, their effects can be mitigated via application of digital filters incorporating slice collection, multiband factor, and repetition time. The implementation of these filters is provided both in interactive online notebooks and an open source denoising tool.

## Introduction

Physiological recordings collected simultaneously during functional magnetic resonance imaging (fMRI) can add valuable information about a participant’s physical state and provide quantitative assessment of psychological phenomena. They offer the opportunity to study relations between the central and autonomic nervous systems that underlie cognition and behavior. For example, physiological arousal has been assessed during fMRI using heart rate, via electrocardiogram (ECG) recordings and skin conductance, via electrodermal activity (EDA) recordings. Inclusion of such measures may enhance interpretation of studies examining decision making (reviewed in (1), typical and disordered affective processing (2–5), pain (6–8), autonomic regulation (9–13). Furthermore, heart rate has known effects on the BOLD signal (14–16) plays a nontrivial role in the emergence of large-scale functional brain networks (17). Thus, measures of heart rate play a role, too, in fMRI denoising (18–20).

Collecting electrophysiological recordings in the MR environment adds MR-induced artifacts to the recordings. Often, the magnitude of these MR artifacts is much larger than that of the phenomena of interest, necessitating additional data cleaning steps before such data can be used to assess psychophysiological phenomena. Single-echo MRI sequences (Figure 1A) that measure the blood-oxygenation level dependent (BOLD) signal are the norm in fMRI research. However, recent advances in MR technology and denoising approaches are prompting researchers to increasingly turn to multi-echo sequences (Figure 1B), which offer better differentiation between BOLD signal and non-neural noise for improved estimates of brain activation (21–23). While these sequences arguably offer better quality fMRI data, they require more complex K-space sampling to collect multiple echoes, which introduces added artifacts to simultaneously collected electrophysiological recordings. Furthermore, adding echoes to an MRI sequence introduces further limitations on the temporal resolution of the sequence, requiring a longer repetition time (TR). Using multiband or simultaneous multi-slice (SMS) excitation allows researchers to reduce the amount of time required to acquire a single volume, by acquiring several slices simultaneously, and effectively minimizing the multi-echo temporal constraints on TR. This is an important consideration for human fMRI studies, in which a study’s power to detect an effect is linearly related to the number of timepoints in a scan (24).

**Figure 1.**
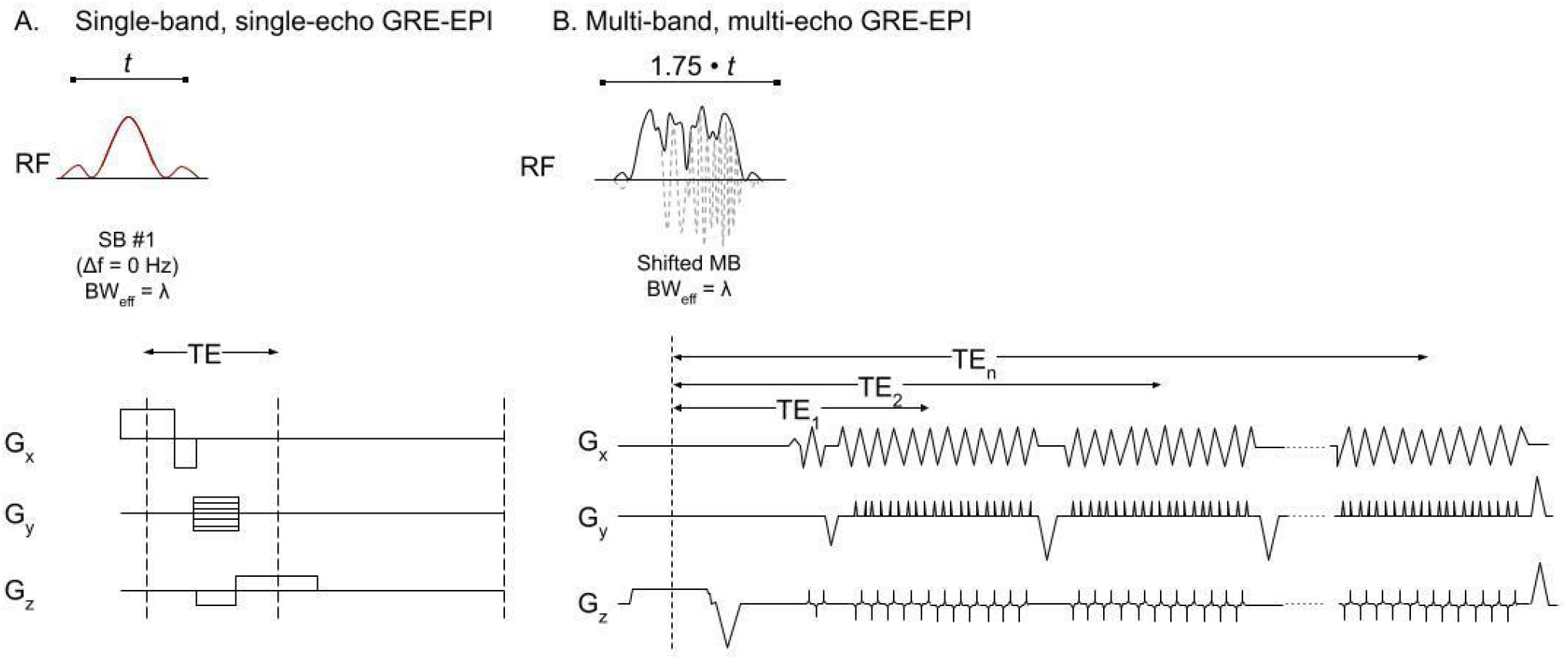
Schematic representation of (A) a single-band, single-echo GRE-EPI pulse sequence and (B) a multi-band, multi-echo GRE-EPI pulse sequence. RF pulse differences **(25)** are highlighted in the first row, while differences in time-varying gradient fields **(26)** are shown in the following three rows. NOTE: This schematic presents a general example of differences between single-band, single-echo sequences and multi-band, multi-echo sequences and is not specific to those sequences described here.

In addition to the technical challenges, neuroimaging researchers who are new to physiological monitoring likely encounter logistical and educational hurdles as well. There are several manufacturers that provide options for physiological monitoring in the MR environment, including BIOPAC Systems, Inc. (Goleta, CA United States), ADInstruments (Sydney, Australia), and the MRI scanner manufacturers themselves. While all provide MR-safe equipment for recording physiological data safely in the scanner, their resources are not widely available. Of these, BIOPAC is a popular choice among neuroimaging researchers and among the only manufacturers to offer recommendations for filtering MR-noise out of concurrently collected electrophysiological data. There is a growing need for transparent, open-source resources for integrating physiological data into neuroimaging research. Several such efforts are currently underway, including NeuroKit (neuropsychology.github.io/neurokit) and the Physiopy tool suite and associated community (github.com/physiopy). However, there remains a gap where open tools and recommendations for filtering noise associated with MR-sequence specifics are concerned.

The present study assessed MR-artifact removal strategies from electrophysiological (i.e., ECG and EDA) data collected during single- and multi-echo multiband EPI scans, focusing on data collected with BIOPAC systems and the manufacturer’s filtering recommendations. Our goals were to (1) compare MR-related noise from single- and multi-echo EPI sequences, (2) assess current filtering recommendations in multiband and multi-echo contexts, and (3) if current filtering recommendations appeared insufficient for removing MR-related artifacts from concurrent electrophysiological data, to redefine these recommendations accordingly. To achieve this, we used ECG and EDA data collected during both single- and multi-echo, single- and multi-band BOLD EPI sequences, from three separate datasets. First, data were Fourier transformed to identify MR-artifact frequencies, then digital filters were applied, and cleaned data were compared across the filtering process, both visually and quantitatively. We anticipated that MR-artifacts would be greatest during multiband, multi-echo EPI sequences and while current filtering recommendations would mitigate MR-artifacts, adaptations may be necessary for multiband and multi-echo pulse sequences. We expected that adaptations to the slice collection frequency would be necessary, to account for slices collected in parallel. Finally, we make these findings openly available both as interactive online code and easy-to-use command-line software that is compatible with current data standards, sharing updated recommendations for removing MR-artifacts from these physiological data in multiband and multi-echo contexts.

## Methods

Data used here come from three separate studies. The first is a larger study of younger and older adults (N = 52: 26 younger (i.e., aged 13-34 years; 9 female), and 26 older (i.e., aged 55-75 years; 9 female) collected at the University of Southern California Dana and David Dornsife Neuroimaging Center (hereafter the Mather dataset; https://doi.org/10.18112/openneuro.ds001242.v.1.0.0; (27–29)) including electrocardiogram (ECG) and electrodermal activity (EDA) data collected during single-band, single-echo BOLD fMRI scans. The second and third are two pilot studies collected at Florida International University’s Center for Imaging Science: one of children that includes ECG and EDA data collected during multiband, single-echo BOLD fMRI scans (N = 2, male, aged 9 years; hereafter the Musser dataset) and one of adults that includes ECG and EDA data collected during multiband, multi-echo BOLD fMRI scans (N = 5, all female, aged 26-39 years; hereafter the DIVA dataset; https://doi.org/10.18112/openneuro.ds002278.v1.0.1). More information about the physiological data and concurrent BOLD fMRI sequences for each study are provided below.

### Physiological recordings

Physiological data for the Mather dataset was acquired using MRI-compatible modules, leads, and electrodes from BIOPAC Systems. Data were acquired using a BIOPAC MP150 system, connected to subject leads. Electrocardiogram (ECG) recordings were collected using radiotranslucent EL508 electrodes with GEL100 and LEAD108 leads, with an ECG100C-MRI amplifier (27).

Physiological data for the Musser and DIVA datasets were acquired using MRI-compatible modules, leads, and electrodes from BIOPAC Systems. Data were acquired using a BIOPAC MP150 system, connected to subject leads by two standard MEC-MRI cables that passed through the MRI patch panel via MRI-RFIF filters and ran without loops to the bore, then parallel with the subject. Electrocardiogram (ECG) recordings were collected using radiotranslucent EL508 electrodes with GEL100 and 15cm long LEAD108B leads, with an ECG100C-MRI amplifier. Electrodes were placed in a 3-lead bipolar monitoring configuration, 6 to 8 inches apart diagonally across the heart from right clavicle to left rib cage, with the ground placed 6 to 8 inches away on the right rib cage. Electrodermal activity (EDA) recordings were collected using radiotranslucent EL509 electrodes with GEL101 and LEAD108B leads, with an EDA100C-MRI amplifier. Leads were placed on the palm of the participant’s non-dominant hand, on the thenar and hypothenar eminences. All physiological data were collected at a rate of 2000 Hz, with ECG and EDA collected concurrently from all participants. Physiological data collection began once participants were loaded on the scanner bed and continued until the scanner bed exited the bore after the scanning session, including several minutes per participant of data collected in the absence of an MR pulse sequence.

### BOLD EPI Sequences

#### Single-band, single-echo BOLD EPI sequence (Mather)

Physiological recordings were acquired in the bore of a whole-body 3-Tesla Siemens MAGNETOM Prisma with a 32-channel head/neck coil during a single-band, single-echo (SBSE) blood-oxygenation-level-dependent (BOLD) echo planar imaging (EPI) sequence (Table 1, Mather). The SBSE sequence used here is a gradient-echo EPI sequence that acquired 41 axial slices using a single echo (TE = 25ms) with TR = 2000 ms, no multiband acceleration, interleaved acquisition, and a 90° flip angle. Participants completed one 6-minute run of a fear conditioning task, and five runs of a spatial detection task that each lasted 5:20.

**Table 1.**
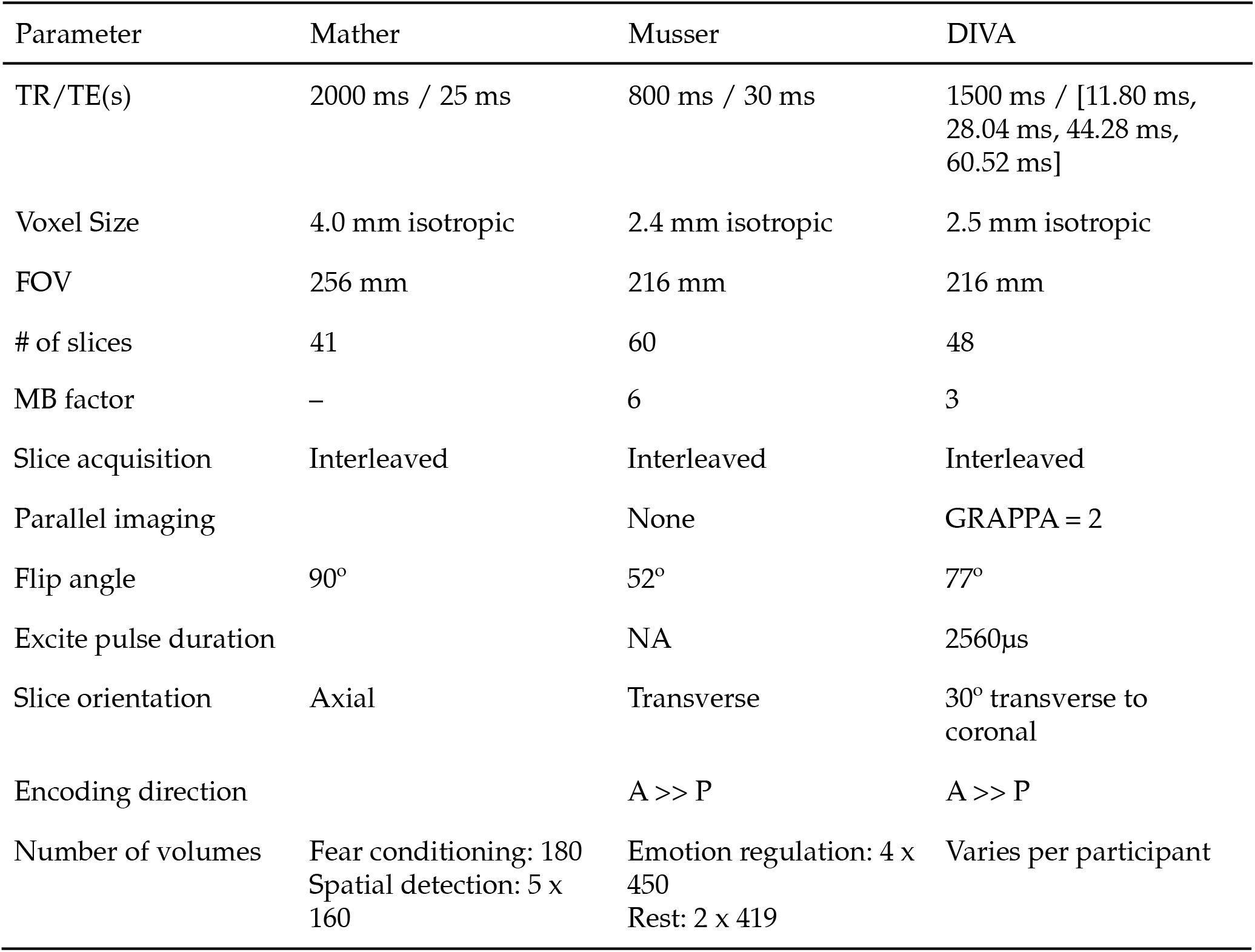
BOLD sequence parameters for each dataset.

#### Multiband, single-echo BOLD EPI sequence (Musser)

Physiological recordings were acquired in the bore of a whole-body 3-Tesla Siemens MAGNETOM Prisma with a 32-channel head/neck coil, during both a multiband, single-echo (MBSE) BOLD EPI sequence (Table 1, Musser). The MBSE sequence used here is the one developed for and used by the Adolescent Brain Cognitive Development (ABCD)^SM^ Study (30). In brief, this sequence acquired 60 transverse slices, with an anterior to posterior phase encoding direction, using a single echo (TE = 30ms) with TR = 800ms, a multiband acceleration factor of 6, interleaved acquisition, in-plane GRAPPA acceleration, and a 52° flip angle. More information about the scan protocols is available with the curated ABCD data via the NIMH Data Archive (NDA; https://abcdstudy.org/scientists/protocols/). Participants completed four runs of an emotion regulation task (31–33) that each lasted 5:35 and two 5-minute runs of rest.

#### Multiband, multi-echo BOLD EPI sequence (DIVA)

Physiological recordings were acquired in the bore of a whole-body 3-Tesla Siemens MAGNETOM Prisma with a 32-channel head/neck coil during a multiband, multi-echo (MBME) BOLD EPI sequence (Table 1, DIVA). The MBME BOLD EPI sequence used here is from the distribution of multi-band accelerated EPI sequences (Moeller et al., 2010) developed by the Center for Magnetic Resonance Research (CMRR) at the University of Minnesota. The MBME GRE-EPI sequence acquired 48 slices at a 30° transverse-to-coronal orientation with anterior-to-posterior phase encoding direction at each of 4 echoes (TE_1_ = 11.80ms, TE_2_ = 28.04ms, TE_3_ = 44.28ms, TE_4_ = 60.52ms) with TR = 1500ms, a multiband acceleration factor of 3, interleaved acquisition, in-plane GRAPPA acceleration, a 77° flip angle, and an excite pulse duration of 2560μs (Figure 1B). Participants completed six runs, 6 to 11 minutes each, of film watching (34), two runs of the same emotion regulation task (35), two runs of a probabilistic selection task, one 6 minutes and the other 9 minutes (36), and two runs of 5 minutes of rest.

The full parameters and fMRI data are available on OpenNeuro.org (https://openneuro.org/datasets/ds002278/versions/1.0.1).

### Software tools

All code used to create and apply these filters is available on GitHub (https://github.com/62442katieb/mbme-physio-denoising), and was written and run using Python 3.7.3. The bioread library (https://github.com/uwmadison-chm/bioread, v. 1.0.4) was used to read in physiological recordings stored in AcqKnowledge format, data were manipulated using Pandas (https://pandas.pydata.org/, v. 1.0.3), digital filters were created and applied using SciPy (https://www.scipy.org, v. 1.9.0; (37)), and fast Fourier transforms implemented in NumPy (https://numpy.org/, v. 1.18.2; (38)).

### Denoising electrocardiogram recordings

Fourier transform was applied to ECG and EDA data collected both in the presence and absence of MR pulse sequences to identify the frequencies of MR-related artifacts. Then, the same for each the ECG and EDA data, we applied digital filters to mitigate the effects of these artifacts on the recordings.

### BIOPAC filtering

First, we applied the manufacturer (i.e., BIOPAC) recommendation for single-band, single-echo sequences: comb band-stop filters at the slice collection frequency and its harmonics up to the Nyquist frequency, and then Fourier transformed the results to assess how these filters mitigated artifacts. This slice collection frequency is defined as:

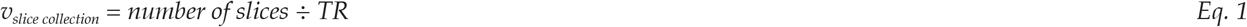

Here, comb band-stop filters were implemented as infinite impulse response (IIR) comb notch filters to account for the fundamental frequency and its harmonics. These filters were then applied to the raw recordings and the resulting filtered signal was Fourier transformed.

### Bottenhorn filtering

Here, we present an update to this calculation, adjusting the slice collection frequency to account for the additional artifacts identified in frequency spectra due to multiband slice acquisition (Eq. 2).

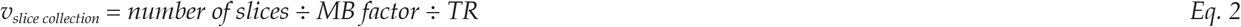

Again, comb band-stop filters were implemented as IIR comb notch filters with this adjusted slice frequency. These filters were then applied to the raw recordings and the resulting filtered signal was Fourier transformed. Further, additional IIR notch filters centered at the TR frequency were implemented and applied to mitigate the effects of additional confounding frequencies present in MBME-ECG recordings.

### Denoising assessments

The quality of ECG data was estimated before and after filtering using two automated approaches: kurtosis and a heuristic-based signal quality index (SQI) (39). Kurtosis has been previously used and evaluated as a signal quality index (SQI) for ECG data (40, 41), such that higher kurtosis is indicative of higher quality. Kurtosis was calculated using Fisher’s definition of kurtosis, as applied by the pandas Python package (v1.0.5) (42, 43). Wilcoxon signed-rank tests were used to compare kurtosis between raw and filtered data. The heuristic quality index (hereafter Zhao heuristic) delineates between “unacceptable”, “barely acceptable”, and “excellent” quality ECG signals by performing a fuzzy comprehensive evaluation of four SQIs: R peak detection, QRS wave power spectrum distribution, kurtosis, and baseline relative power. The Zhao heuristic was computed by the ecg_quality function from NeuroKit2 (https://neuropsychology.github.io/NeuroKit/; (44)), which calculates the four SQIs and synthesizes them into a single quality estimate as described in detail by Zhao and Zhang (2018) (39). Chi-squared tests were used to compare heuristic frequencies between filtering approaches.

Finally, physiological data were compared across steps and to data collected in the absence of MR pulse sequences, using magnitude squared coherence to assess linear dependence across the frequency band in which physiologically-relevant signals were found: 0.5 - 50Hz for ECG and <0.5Hz for EDA. Magnitude squared coherence (Cxy) measures the linear dependence between two vectors (x, y) and can be used to assess the similarity of frequencies between two signals. Here, we use Welch’s method ((45); Eq. 3), as implemented in SciPy (v1.9.0):

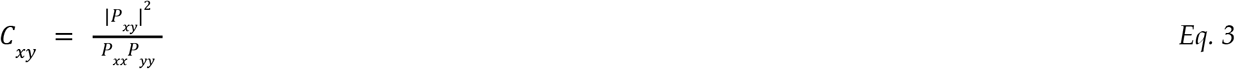

Where *P_xy_* is the cross spectral density estimate of signals x and y; *P_xx_* and *P_yy_* are the power spectral densities of x and y, respectively.

This allowed a direct comparison of frequency spectra between (i) filtering approaches, between (ii) filtered recordings and unfiltered recordings, and between (iii) filtered recordings and recordings in the absence of EPI sequences. In this case, these comparisons show which frequencies have been removed between filtering steps (low C_*xy*_) and which frequencies remain (high C_xy_).

## Results

### Frequencies of MR-related artifacts

Comparing Fourier transforms to ECG recordings in both the absence (Figure 2, first row) and presence (Figure 2, second row) of single-band single-echo (Figure 2, left column), multiband multi-echo (Figure 2, center column) and multiband multi-echo (Figure 2, right column) BOLD EPI sequences (hereafter SBSE-ECG, MBSE-ECG, and MBME-ECG, respectively) revealed MR-related noise in frequencies corresponding to the TR and slice acquisition. This noise demonstrated greater spectral power in recordings collected during multi-echo sequences than during single-echo sequences (Figure 2D vs. 2L). The presence and relative impacts of this noise are visually apparent in the difference between recordings collected before and during the two BOLD EPI sequences in Figure 2 (MBSE-ECG in 2A vs. 2C; MBME-ECG in 2I vs. 2K) and evidenced by greater power in the frequencies corresponding with TR and slice acquisition than with biologically-relevant signals (MBSE-ECG in 2B vs. 2D; MBME-ECG, 2J vs. 2L). These artifacts occur at frequencies equal to (a) the multiband *slice frequency* which is equal to the number of slices divided by the multiband factor per TR (indicated by circular, blue-green markers), (b) the *TR frequency* (indicated by triangular, pink markers), and the harmonics of these frequencies. Furthermore, the power of these confounding signals was much greater than in MBSE-ECG recordings and caused greater corruption of the ECG signal (Figure 2C vs 2K).

**Figure 2.**
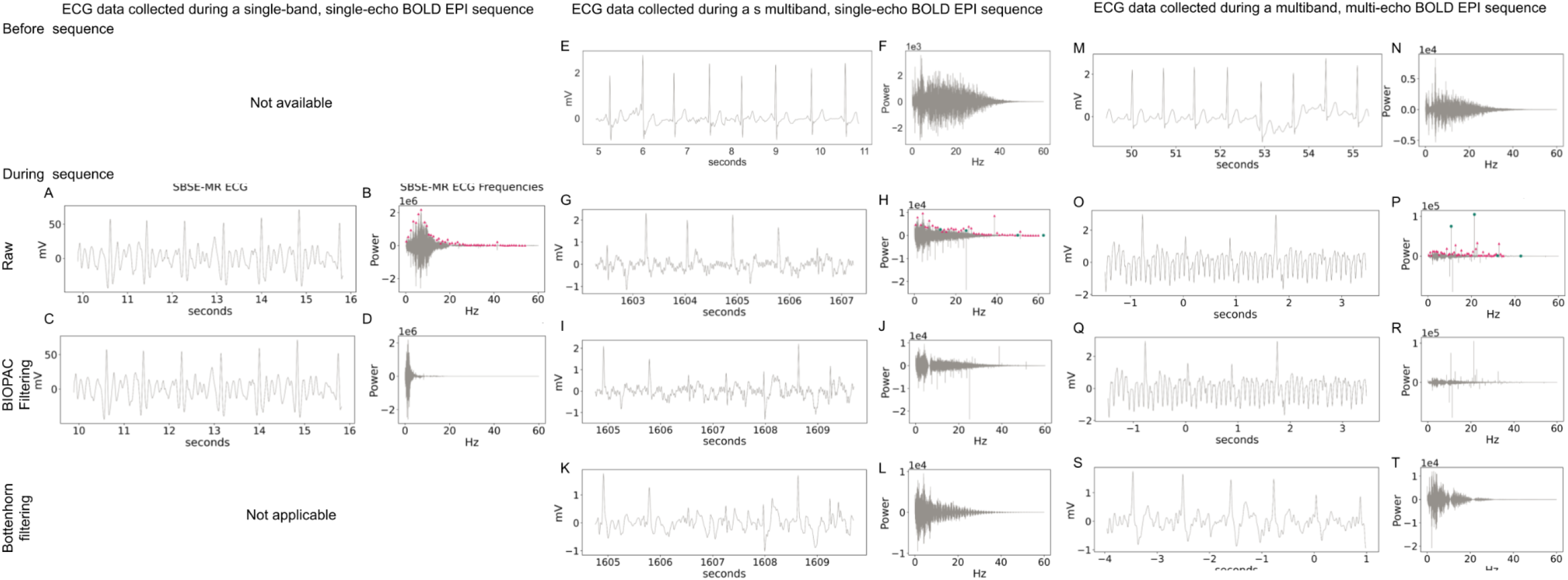
Electrocardiogram recordings through the denoising process. *ECG during single-band, single-echo (SBSE) BOLD EPI sequence:* (A) Six seconds of ECG recording during a single-band, single-echo BOLD EPI sequence and (B) the Fourier transform of that recording. (C) Six seconds of ECG recordings during a single-band, single-echo BOLD EPI sequence and (D) the Fourier transform of that recording, demonstrating sequence-related artifacts at the slice frequency (green circles) and TR frequency (pink triangles), and their harmonics. *ECG during multiband, single-echo (MBSE) BOLD EPI sequence:* (E) Six seconds of ECG recordings before the EPI sequence started, after the participant was moved into the scanner bore and (F) the associated power spectrum, which shows remaining MR-related artifacts after filtering. (G) Six seconds of ECG recordings during a multiband, single-echo BOLD EPI sequence and (H) the Fourier transform of that recording, demonstrating sequence-related artifacts at the slice frequency (green circles) and TR frequency (pink triangles), and their harmonics (I) The same 6 seconds of ECG recordings, following application of IIR notch filters to remove the MR-related artifacts, per manufacturer recommendations and (J) the associated power spectrum, which shows remaining MR-related artifacts after filtering. (K) The same 6 seconds, following application of IIR notch filters updated for multiband acquisition and (L) the associated power spectrum, displaying mitigated artifacts, but not complete removal. *ECG during multiband, multi-echo (MBME) BOLD EPI sequence:* (M) 6 seconds of ECG recordings from an individual in the MR environment prior to scanning with a BOLD EPI sequence and (N) the power spectrum of that pre-EPI recording. (O) 6 seconds of ECG recordings from the same individual and scanning session during a multiband, multi-echo BOLD EPI sequence data and (P) the power spectrum of that ECG recording, demonstrating MR-related artifacts at the slice frequency (green circles) and TR frequency (pink triangles), and their harmonics. (Q) The same 6 seconds of ECG recording, following the application of BIOPAC-recommended filters and (R) the Fourier transform of that recording, displaying the remaining MR-related artifacts. (S) The same 6 seconds of ECG recording, following the application of IIR notch filters at the slice and TR frequencies and (T) the Fourier transform of that cleaned recording, displaying the relative absence of MR-related artifacts. NOTE: Power spectra (i.e., plots of signal Fourier transforms) y-axes are log10-scaled.

### Denoising electrocardiogram recordings

BIOPAC-recommended filtering applied to SBSE-, MBSE-, and MBME-ECG recordings, via IIR notch filters, resulted in an incomplete mitigation of MR-related artifacts, which are still clearly present in the frequency spectra (Figure 2F, N). However, the adjusted slice frequency, and in the case of MB-ECG data at TR frequency, filters mitigated the effects of the identified MR scanner noise in both MBSE- and MBME-ECG recordings, per visual inspection of the power spectra (Figure 2H, P).

### Denoising electrodermal activity recordings

Prior research on EDA recordings collected during single-band, single-echo BOLD sequences has shown minimal MR-related artifacts in EDA data (46). As such, EDA recordings collected simultaneously with fMRI data do not typically require MR-specific denoising. The EDA recordings acquired during a single-band, single-echo BOLD EPI sequence (hereafter SBSE-EDA) included here (Figure 3A, B) support this idea, showing minimal MR-related artifacts and, instead, induced artifacts following filtering (Figure 3C, D). On the other hand, a Fourier transform of EDA recordings acquired during the multiband, single-echo BOLD EPI sequence (hereafter MBSE-EDA) in question revealed noise in sequence-specific frequency bands (Figure 3D, L) corresponding to the harmonics of the TR frequency (pink triangles) and, to a lesser extent, the slice collection frequency (green circles).

**Figure 3.**
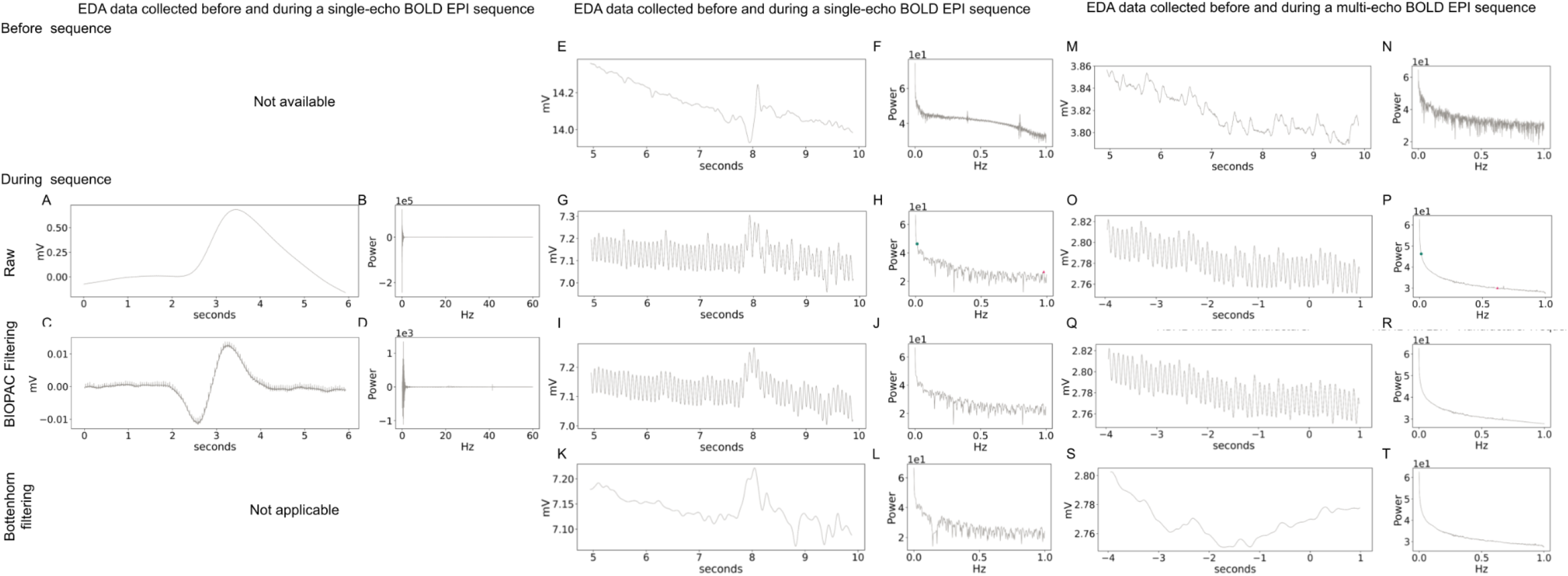
Electrodermal activity recordings through the denoising process. *EDA during single-band, single-echo (SBSE) BOLD EPI sequence:* (A) Six seconds of EDA recording during a single-band, single-echo BOLD EPI sequence and (B) the Fourier transform of that recording. (C) Six seconds of EDA recordings during a single-band, single-echo BOLD EPI sequence and (D) the Fourier transform of that recording, demonstrating sequence-related artifacts at the slice frequency (green circles) and TR frequency (pink triangles), and their harmonics. *EDA during multiband, single-echo (MBSE) BOLD EPI sequence:* (E) Six seconds of EDA recordings before the EPI sequence started, after the participant was moved into the scanner bore and (F) the associated power spectrum, which shows remaining MR-related artifacts after filtering. (G) Six seconds of EDA recordings during a multiband, single-echo BOLD EPI sequence and (H) the Fourier transform of that recording. (I) The same 6 seconds of EDA recordings, following application of IIR notch filters to remove the MR-related artifacts, per manufacturer recommendations and (J) the associated power spectrum, which shows remaining MR-related artifacts after filtering. (K) The same 6 seconds, following application of IIR notch filters updated for multiband acquisition and (L) the associated power spectrum, displaying mitigated artifacts, but not complete removal. *EDA during multiband, multi-echo (MBME) BOLD EPI sequence:* (M) 6 seconds of EDA recordings from an individual in the MR environment prior to scanning with a BOLD EPI sequence and (N) the power spectrum of that pre-EPI recording. (O) 6 seconds of EDA recordings from the same individual and scanning session during a multiband, multi-echo BOLD EPI sequence data and (P) the power spectrum of that EDA recordings. (Q) The same 6 seconds of EDA recording, following the application of BIOPAC-recommended filters and (R) the Fourier transform of that recording, displaying the remaining MR-related artifacts. (S) The same 6 seconds of EDA recording, following the application of IIR notch filters at the slice and TR frequencies and (T) the Fourier transform of that cleaned recording, displaying the relative absence of MR-related artifacts. NOTE: Power spectra (i.e., plots of signal Fourier transforms) y-axes are log10-scaled.

### Quantitative assessments of denoised data

Filtering approaches differentially impacted the quality of ECG recordings across the three datasets included here. BIOPAC-recommended filtering did not significantly improve ECG signal quality across the Mather (Table 2, SBSE-ECG), Musser (Table 2, MBSE-ECG) or DIVA (Table 2, MBME-ECG) datasets, according to any of the included metrics. The updated filtering recommendations presented here (Table 2, Bottenhorn) significantly improved ECG signal quality in MBSE- and MBME-ECG data compared with the signal filtered per BIOPAC-recommendations, but not compared with the raw signal. However, visual inspection of the example signals in (Figure 2) across filtering approaches show a mitigation of MR-associated noise and, in the case of MBME-ECG data, increased R-peak amplitude compared with the surrounding noise. Thus, while the updated filtering recommendations might not objectively improve signal quality, they might still benefit researchers interested in computing heart rate or variability therein. For distributions of each quality index and heart rate estimates across datasets and filtering approaches, see Supplementary Figure 1.

**Table 2.** Signal quality across filtering approaches, averaged across participants, runs, and sessions.

|  | Raw | BIOPAC | Bottenhorn | Raw < BIOPAC | Raw < Bottenhorn | BIOPAC < Bottenhorn |
| --- | --- | --- | --- | --- | --- | --- |
| SBSE-ECG |  |  |  |  |  |  |
| Kurtosis | 8.14 ± 17.34 | 8.07 ± 17.07 | – | p = 0.99 | – | – |
| Zhao | 44 / 162 / 97 | 42 / 164 / 97 | – | p = 0.94 | – | – |
| MBSE-ECG |  |  |  |  |  |  |
| Kurtosis | 12.75 ± 10.13 | 10.75 ± 10.00 | 12.79 ± 12.4 | p = 1.0 | p = 0.70 | **p = 0.0007** |
| Zhao | 0 / 0 / 12 | 1 / 7 / 4 | 0 / 3 / 9 | – | – | – |
| MBME-ECG |  |  |  |  |  |  |
| Kurtosis | 2.07 ± 1.99<br>(2.08 ± 2.02) | 1.68 ± 2.09<br>(1.69 ± 2.12) | 6.04 ± 22.85*<br>(2.11 ± 2.07) | p = 1.0<br>(p = 1.0) | p = 0.78<br>(p = 0.08) | **p = 0.007**<br>**(p = 0.025)** |
| Zhao | 0 / 123 / 13<br>(0 / 119 / 13) | 0 / 123 / 13<br>(0 / 119 / 13) | 0 / 116 / 20<br>(0 / 116 / 16) | – | p = 0.09<br>(p = 0.38) | p = 0.09<br>(p = 0.42) |

Note: Zhao heuristics are represented as “unacceptable / barely acceptable / excellent”. Bold indicates significantly greater quality index across participants, runs, and sessions (where applicable) at $\alpha < 0.05$ , per 1-tailed Wilcoxon signed-rank test. \*In the MBME-ECG data one participant’s last four scans demonstrated outlier kurtosis values by an order of magnitude after Bottenhorn filtering. Values in parentheses represent the metrics and comparison $p$ -values after
removing those runs from analysis.

Magnitude squared coherence (i.e., linear dependence, or signal similarity across frequencies) of ECG recordings across the filtering approaches described above demonstrated the efficacy of applied notch filters in removing the targeted frequencies from ECG recordings. Linear dependence between ECG recordings collected in the absence of either a single- or multi-echo multiband BOLD EPI sequence and recordings collected during those sequences was near zero across biologically-relevant frequencies. On the other hand, magnitude squared coherence of EDA recordings collected before and during BOLD EPI sequences, and across filtering approaches indicate greater linear dependence between signals collected in the absence of BOLD EPI sequence and those collected during multi-echo than during single-echo BOLD EPI sequence. Across raw and filtered SBSE-, MBSE-, and MBME-EDA data, magnitude squared coherence demonstrated the efficacy of each filter in removing the desired frequencies from the signals. Overall, this supports the claim in prior research that the impacts of MR-related artifacts on simultaneously-collected EDA recordings are minimal, although not nonexistent.

### Research Products

The workflows used to clean these data and create the associated figures are available as a command-line Python script and in interactive Jupyter Notebooks available at: https://github.com/62442katieb/mbme-physio-denoising/.

These notebooks are additionally available, interactively, at: https://mybinder.org/v2/gh/62442katieb/mbme-physio-denoising/binder-live.

## Discussion

Here, we assessed the confounding influence of both single- and multiband, single-echo and multi-echo BOLD MRI sequences on simultaneously acquired peripheral physiological recordings (i.e., ECG and EDA). These artifacts were demonstrated in recordings collected over the course of several MRI scans, comparing those of a SBSE BOLD EPI scan (i.e., from the Mather dataset) with those of a MBSE BOLD EPI sequence with a multiband factor of 6 (i.e., from the Musser dataset) and a MBME BOLD EPI sequence that acquired 4 volumes per RF excitation with a multiband factor of 3 (i.e., from the DIVA dataset). Two fundamental confounding frequencies were identified, corresponding with the slice frequency and the repetition time of the MRI sequence, with notably greater power in the MBSE and MBME sequences, compared with the SBSE sequence. Applying a series of notch filters centered at frequencies corresponding to the sequence’s TR and slice collection frequency, approximating a comb band-stop filter (per manufacturer (i.e., BIOPAC) recommendations) provided marked decrease of confounding signals. Based on this, we present an updated set of recommendations for mitigation of pulse sequence-related artifacts in ECG and EDA recordings collected during multiband BOLD MRI scans. These recommendations make it easier for researchers to include physiological recordings during functional MRI studies that capitalize on the improved temporal signal-to-noise ratio (tSNR) of multi-echo pulse sequences and the improvements to temporal resolution made possible by simultaneous multi-slice acquisition. While we did not test these recommendations across a range of pulse sequences with different numbers of echoes and multiband factors, it is likely that our recommendations will generalize across MBME BOLD EPI sequences due to the linear relationship between confounding frequency bands and the sequence’s TR and multiband factor.

Building on prior research, we found MRI sequence artifacts in simultaneously collected ECG recordings in frequency bands correspond to the number of asynchronously-collected slices (i.e., the number of slices divided by the multiband factor). Furthermore, these confounding frequencies were of greater power in recordings collected during multi-echo BOLD EPI scans than during single-echo. The impact of these corrupting frequencies can be removed with a series of IIR notch filters corresponding to the TR and slice collection frequencies and their harmonics up to the Nyquist frequency. This approach imparts greater increases in R-peak discriminability, though not necessarily objective signal quality indices, to data collected during multiband, single-echo EPI scans than during multiband, multi-echo EPI scans. Further processing is needed in order to distinguish moment-to-moment heart rate and data derived therefrom, but the data have been largely cleaned of the confounding MR-related artifacts.

Contrary to prior research, we found MRI sequence artifacts in simultaneously collected EDA recordings, both during multiband single- and multi-echo BOLD EPI sequences, but not during a single-band, single-echo BOLD EPI sequence. These artifacts corresponded with the TR frequency, likely related to the transmission of RF excitation pulses, and the gradient pulses during slice collection. However, the relative power of these confounding frequencies did not differ between MBSE-EDA and MBME-EDA, in contrast to those in the ECG signals (see Figure 2). Here, we demonstrate that these artifacts can be removed in the same manner as from MBSE- and MBME-ECG recordings with a IIR comb notch filter that removes a frequency band and its harmonics up to the Nyquist frequency. However, given the relatively low frequency of biologically relevant, phasic EDA signals, low-pass filtering may provide a simpler approach for commensurate improvements in signal quality.

Confounding frequencies corresponding with the TR of the sequence were detected via comparison of the power spectra of ECG and EDA recordings before and during a MBME BOLD EPI sequence, to a lesser extent during a MBSE BOLD EPI sequence, and not at all during a SBSE BOLD EPI sequence. This frequency is not often mentioned in the simultaneous physiology-fMRI literature as RF pulse-related artifacts are either of a much lesser amplitude than other MR-artifacts or they are filtered out entirely by MRI-specific amplifiers (see Figure 1 for comparison) (47, 48). However, the RF excitation that precedes slice collection in multiband MRI pulse sequences has a higher amplitude and/or greater total power than that of a single-band sequence. This increased power may explain why the artifact is seen here, in frequency bands corresponding with the sequence TR, but not usually seen in physiological recordings collected simultaneously with single-band BOLD sequences and is not mentioned in prior simultaneous ECG- or EDA-fMRI research or the associated manufacturer recommendations for signal cleaning.

The confounding fundamental frequency identified here corresponds with the slice collection or slice repetition frequency (48). This artifact is more commonly seen in ECG recordings collected during fMRI scans, though not in EDA recordings (46). The literature on simultaneous EEG-fMRI acquisition and data cleaning suggests that the magnitude of artifacts due to electromotive force caused by time-varying magnetic field gradients during slice acquisition far surpasses that of the RF excitation pulse (48). While this artifact is seen in physiological recordings acquired during single-band BOLD sequences, as well, the power of the harmonics of this confounding frequency are much greater in data collected during multiband BOLD sequences. Although slice collection in multi-echo GRE-EPI sequences is more prolonged over the course of a timepoint of data acquisition, due to the acquisition of multiple volumes of data per RF excitation pulse, the duration of slice collection is short (<75ms) compared to the repetition time (1500ms). As such, the confounding frequency associated with time-varying gradients is centered on the slice frequency (slices / MB factor / TR) and confounding frequencies associated with individual echoes were not observed. Notch filters centered at the slice frequency and its harmonics sufficiently removed the artifact caused by shifting gradient fields.

### Limitations and Considerations

The temporal resolutions of each electrophysiological (1000 - 10,000Hz) and fMRI (0.5 - 1.5Hz) data complicate psychophysiological analyses. First, in relating physiological processes to BOLD signal fluctuations, accounting for differences in the timing of individual slice collection, typically performed in the beginning of fMRI preprocessing (49–51), becomes crucial. Second, physiological data should be downsampled for such investigations. fMRI data are collected with a TR between 500ms and 3s and while multiband acquisition can shorten TRs, multi-echo acquisition often lengthens TRs. When TRs exceed 2 seconds, the temporal resolution of fMRI data becomes low enough to induce aliasing in biologically-relevant frequency bands of physiological data downsampled to match. Researchers should proceed with caution when this is the case. Further, future work should consider the differential effects of MR-related artifacts across the spectrum of heart rate and examine potential differences in filtering efficacy, signal quality, and potential impacts on fMRI results, especially given known effects of heart rate on BOLD signal.

On another note, researchers collecting data regarding heart rate and cardiac pulsations can avoid MR artifacts entirely by using a photoplethysmograph (PPG), which collects optical instead of electrical measurements. Both ECG and PPG are used by the neuroimaging community and in a recent mega-analysis of cardiac function and cortical thickness that pooled structural MRI and heart rate data from 20 research groups, half of the groups used PPG to estimate heart rate; the other half, ECG (52). Both ECG and PPG have strengths and weaknesses, however, and the choice between the two depends on investigators’ intended use of the data. Cardiac data can be useful in mitigating the effect of cardiac pulsations on the BOLD signal itself (i.e., for denoising fMRI data) or to investigate the neural correlates of cardiac function. For denoising concurrently collected fMRI data using approaches like RETROICOR (53) or DRIFTER (54), cardiac pulse estimates from either ECG or PPG can be used. In this case, PPG has the advantage as its sensor is easier to place and its signal is less susceptible to MR-induced artifacts (20). On the other hand, data from ECG and PPG are not equally well-suited to some assessments of the neural bases of cardiac function. While they provide nearly equivalent estimates of heart rate, PPG does not provide reliable estimates of heart rate variability (55–58). Finally, while placement of a PPG sensor on the finger or foot is easier, quicker, and less invasive than that of electrodes on the thorax, PPG is susceptible to motion-induced noise (59), which is greater in some populations than others. Altogether, choice of PPG or ECG for cardiac monitoring should depend on the data’s intended use and the concomitant MR sequence.

Finally, the findings presented here only reflect data collected on BIOPAC physiological monitoring systems in SIEMENS MRI scanners. Future work should ascertain the efficacy of such filtering approaches on data collected in other environments and by other systems.

## Conclusions

While MBSE and MBME pulse sequences introduce more complicated artifacts into simultaneously acquired electrophysiological recordings than can be addressed with current manufacturer recommendations, the data presented here suggest that these artifacts are predictable and their effects can be greatly mitigated with notch filters centered at their fundamental frequencies and harmonics. By targeting the slice acquisition frequency, updated to account for multiband factor, and, especially in the case of MBME-simultaneous recordings, TR frequency, researchers should be able to remove significant MR-related artifacts from ECG and EDA data collected during fMRI scans. Recommendations such as those demonstrated here allow researchers to capitalize on the improved SNR afforded by MBSE and MBME BOLD sequences, while including rich information concerning a participant’s peripheral, visceral state.

## Supplemental Material

Supplemental material for this article is available online.

