## Supplementary Material for "Denoising physiological data collected during multi-band, multi-echo EPI sequences"

This document includes:

- Supplemental Figures 1 to 5
- Supplemental References

*Corresponding Author*: **Dr. Katherine Bottenhorn**, Postdoctoral scholar, Department of Population and Public Health Sciences, University of Southern California, Los Angeles, CA, USA |

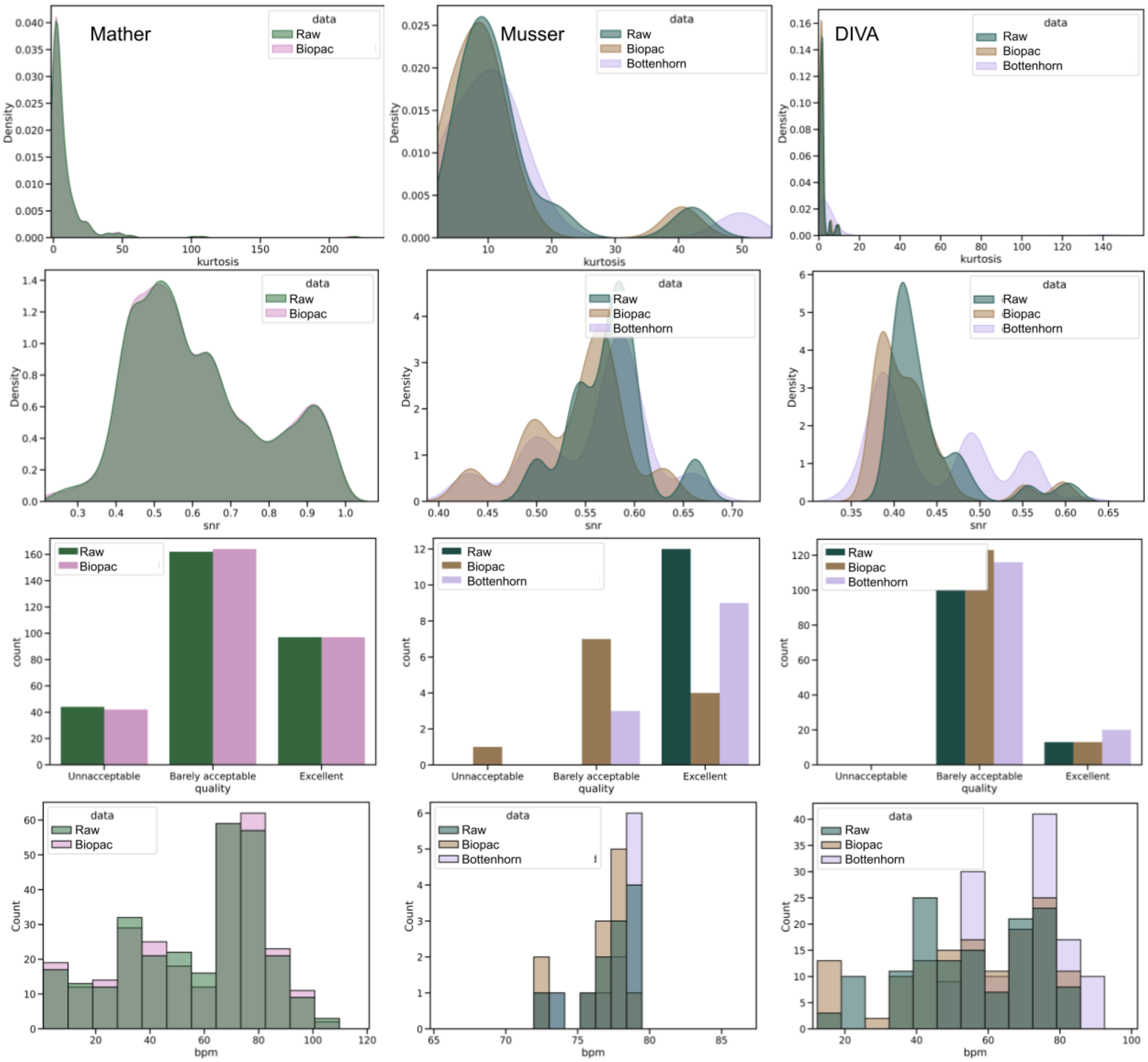


***Supplementary Figure 1.*** Distributions of signal quality indices (top 3 rows) and heart rate (bottom row) from ECG data across datasets (columns) and filtering approaches (colors).

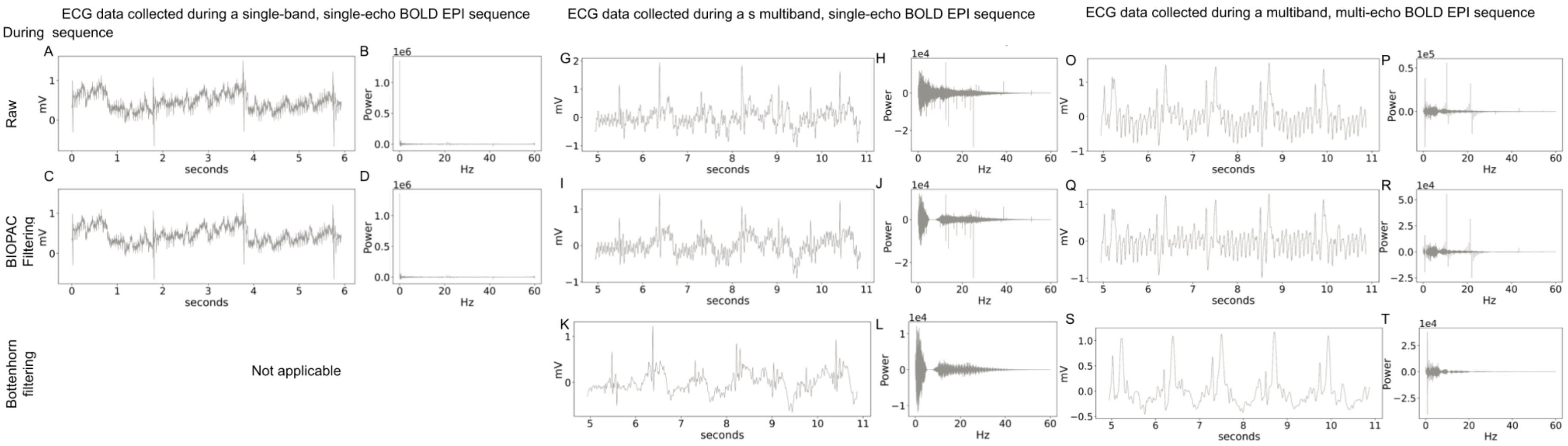


*Supplementary Figure 2.* Examples of ECG signal and the associated power spectra, across filtering approaches, from participants with heart rates that were lower than average for the sample, from each dataset.

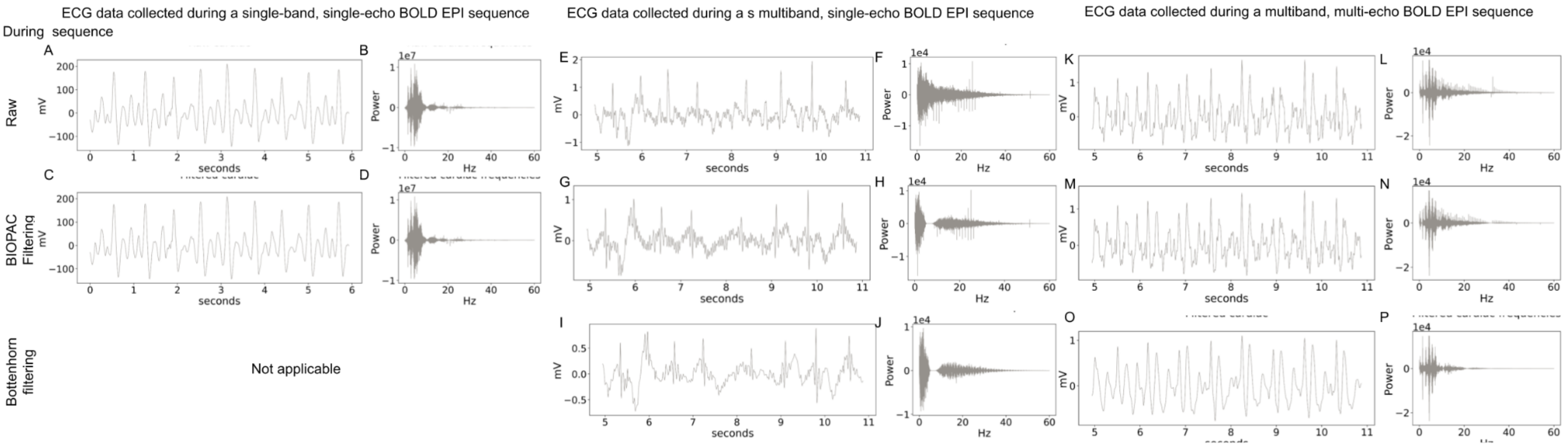


*Supplementary Figure 3.* Examples of ECG signal and the associated power spectra, across filtering approaches, from participants with heart rates that were higher than average for the sample, from each dataset

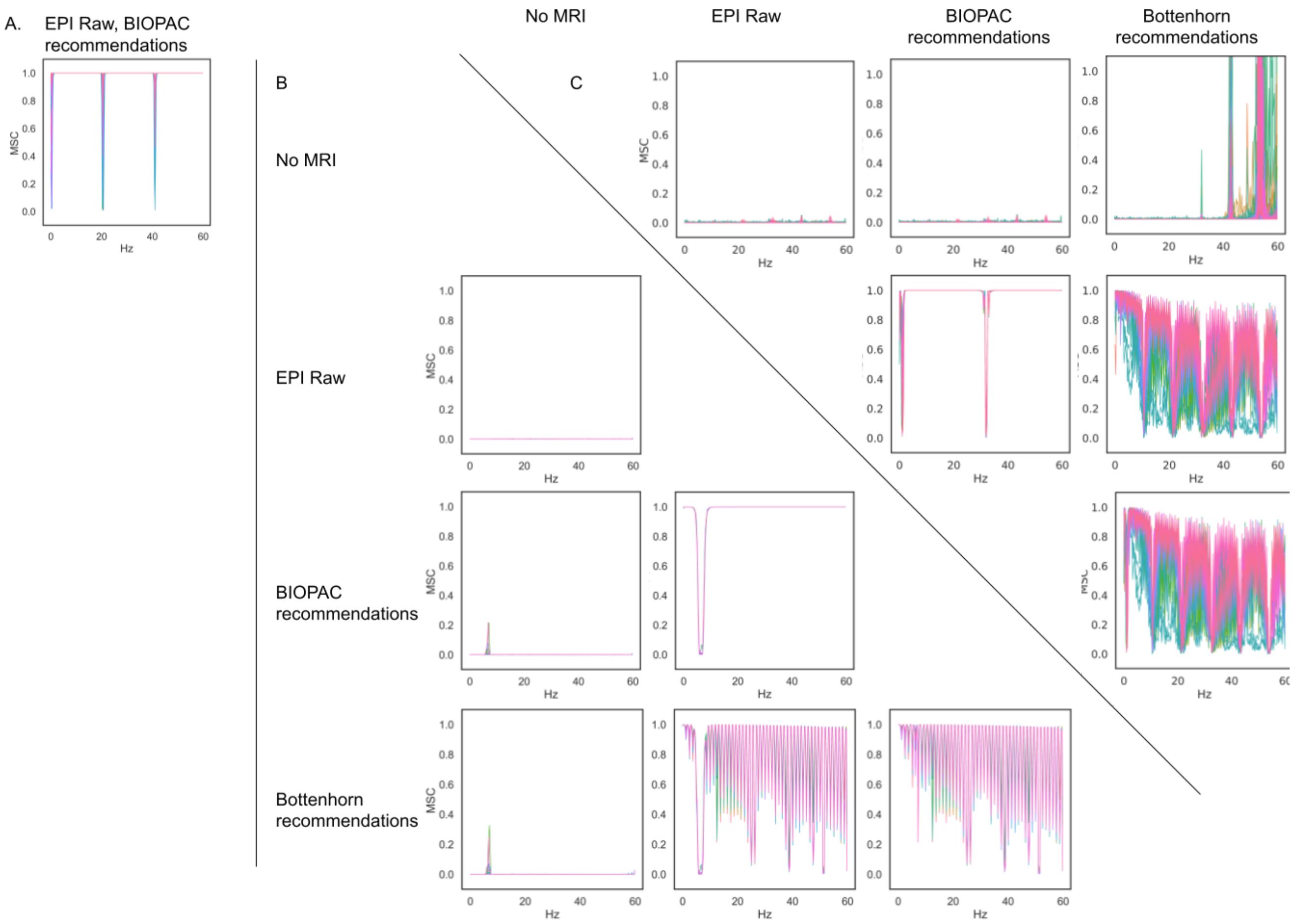


***Supplementary Figure 4.*** Magnitude squared coherence (MSC) between raw and cleaned ECG recordings. (A) Comparison between SBSE-ECG recordings (raw, filtered per manufacturer recommendations). (B) Comparisons between MBSE-ECG recordings are shown below the diagonal; (C) comparisons between MBME-ECG recordings, above the diagonal. Colored lines represent each subject; in the case of DIVA, each session per subject.

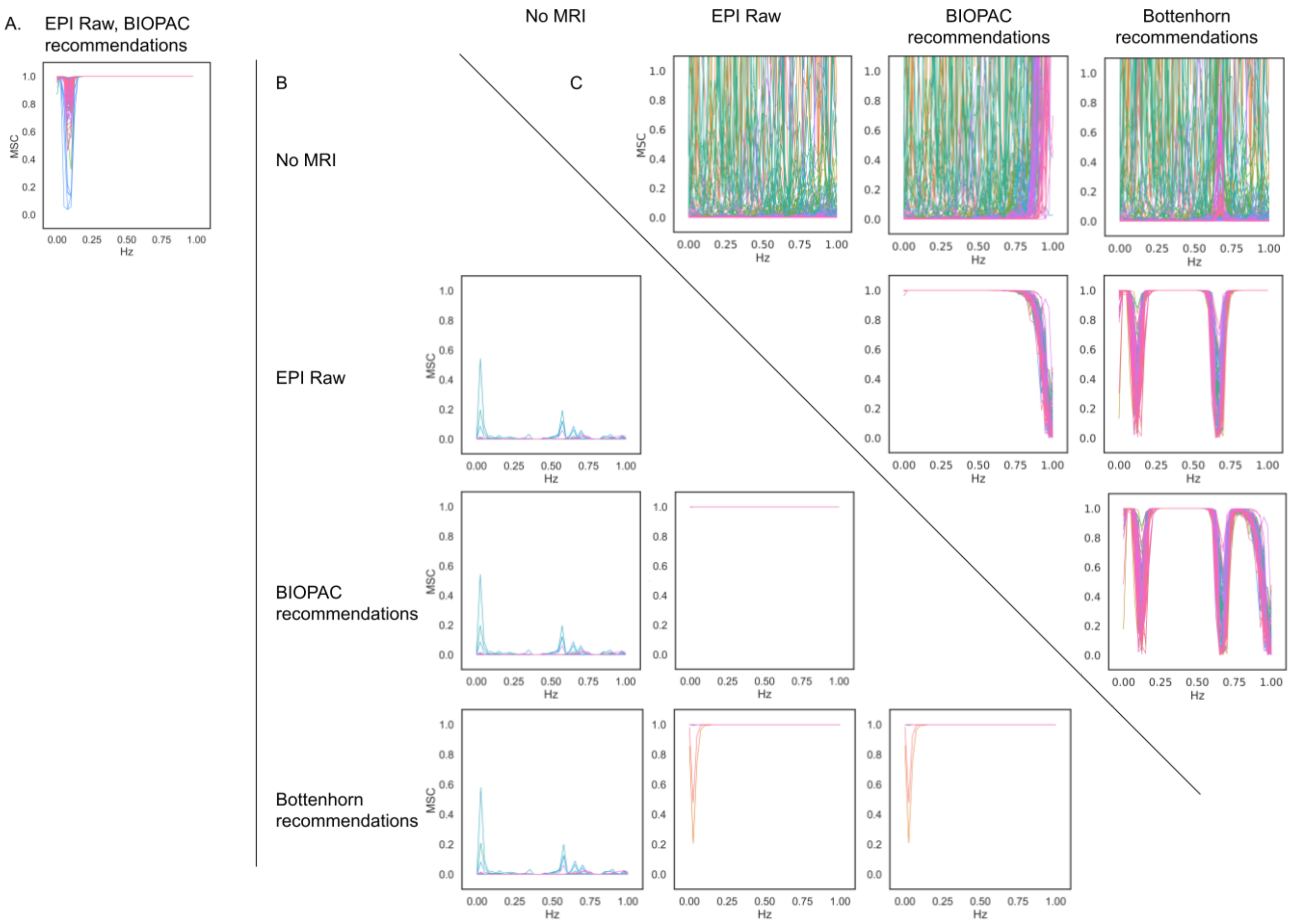


***Supplementary Figure 5.*** Magnitude squared coherence (MSC) between raw and cleaned EDA recordings. (A) Comparison between SBSE-EDA recordings (raw, filtered per manufacturer recommendations). (B) Comparisons between MBSE-EDA recordings are shown below the diagonal; (C) comparisons between MBME-EDA recordings, above the diagonal. Colored lines represent each subject; in the case of DIVA, each session per subject.
